# Cytosolic water potential as a mechanistic driver of leaf airspace unsaturation and non-stomatal control of transpiration

**DOI:** 10.1101/2025.07.08.663815

**Authors:** Diego A. Márquez, Lucas A. Cernusak, Florian A. Busch, Graham D. Farquhar

**Affiliations:** School of Biosciences and Birmingham Institute of Forest Research, University of Birmingham, Edgbaston, Birmingham B15 2TT, UK; Plant Sciences, Research School of Biology, Australian National University, Canberra, ACT 2601, Australia; College of Science and Engineering, James Cook University, Cairns, Queensland 4878, Australia

**Author notes:** Corresponding author: Diego A. Márquez. Authors BlueSky: @dmarquez.bsky.social (D.A.M.); @lucascernusak.bsky.social (L.A.C.); @florianabusch.bsky.social (F.A.B.).

**Keywords:** humidity gradients, water potential, mesophyll cell wall, gas exchange

## Abstract

Mesophyll cells exhibit a previously underappreciated capacity to regulate water loss via low plasma membrane conductance (*L*_p_), offering a non-stomatal mechanism for transpiration control. However, the structural basis and regulation of *L*_p_ remain poorly understood, limiting its integration into predictive models. In this study, we show that *L*_p_ responds dynamically to changes in cytosolic water potential (*ψ*_cy_), decreasing as *ψ*_cy_ approaches the turgor loss point. This identifies *ψ*_cy_ as the primary physiological signal regulating *L*_p_. We introduce a predictive, physiologically grounded model linking *L*_p_ to *ψ*_cy_. Our model establishes a mechanistic connection between internal water status, leaf hydraulics, substomatal cavity unsaturation, and gas exchange. This framework opens new avenues for understanding and modelling plant water use under stress.

## 2 Main

Improving water use efficiency (WUE), the relation between carbon gain and water loss, remains a central goal for agriculture under increasing climatic pressure. While stomatal regulation has long been viewed as the primary control over transpiration^1–4^, recent work has shown that a substantial portion of water loss is regulated independently of stomatal aperture^5–9^. This non-stomatal control—arising from a decrease in conductance to water flow across the mesophyll cell membrane (*L*_p_)^7,9^—has been shown to reduce water loss without impacting carbon assimilation under evaporative stress. In C_4_ species, non-stomatal regulation has been shown to account for over 90% of the capacity to maintain high assimilation and WUE under moderate to high vapour pressure deficit (VPD, ∼2–3 kPa), and contributes around 30% in C_3_ species under similar conditions^9^. These findings highlight the physiological relevance and potential of non-stomatal control for improving crop performance in water-limited environments.

The non-stomatal control of transpiration (i.e., *L*_p_) entails water loss being regulated not just by the stomatal aperture but also by how readily water exits cells into the evaporating surface. This kind of control has been hypothesised for decades^5,10–14^, but compelling evidence of *L*_p_ existence and function has appeared in recent years^5,7,9^, marking a major conceptual shift in plant physiology. However, many fundamental aspects of *L*_p_ remain unclear.

In practice, traditional gas exchange models assume that the airspace inside the leaf (*w*_i_) is saturated with water vapour (*w*_i_/*w*_sat_ = 1), meaning the vapour pressure at the site of evaporation equals the saturation vapour pressure at leaf temperature^1^. However, *w*_i_ can fall below saturation, leading to a measurable vapour pressure deficit within the leaf^5,7^. This phenomenon, termed substomatal cavity unsaturation, implies the existence of a low *L*_p_ as it cannot be explained by stomata alone.

Previous studies have established the presence of *L*_p_ by showing large water potential gradients between the cytosol and the cell wall at a single time point^7^, by inferring its effect from the evolution of substomatal unsaturation^5,9^, or by inferring apoplastic water potential using fluorescent nanoreporters^15,16^. To the best of our knowledge, before this study, there was no direct, point-by-point evidence showing how *L*_p_ decrease dynamically and to what cues it responds. Moreover, while the endpoint difference between the cell wall and cytosolic water potential has been established^7^, the continuous relationship between *L*_p_ and cytosolic water potential (*ψ*_cy_) has not been quantified. This limits our ability to understand or predict how plants adjust this internal conductance under changing environmental conditions.

Importantly, *L*_p_ provides a mechanistic link between plant hydraulics and gas exchange. As water exits the cytosol (transpiration rate), diffusing through the plasma membrane, *L*_p_ determines the water potential of the cell wall (*ψ*_wall_), and consequently the water vapour content in the substomatal cavity (*w*_i_)^7,9,14^. Ultimately, *L*_p_ plays a critical role in determining the vapour pressure gradient driving transpiration (*w*_i_ – *w*_a_). In this study, we address the lack of a mathematical framework describing how *L*_p_ is modulated during transpiration, a gap that stems from the limited understanding of its regulation^14,17,18^. Without such a framework, integrating non-stomatal control into models of transpiration and its links to water use and carbon assimilation remains challenging^19^.

A central unresolved question is whether *L*_p_ is triggered by external drivers like vapour pressure deficit (VPD), transpiration fluxes or by internal signals such as *ψ*_cy_. Identifying the physiological signal that drives *L*_p_ is therefore a necessary step towards building a mechanistic model of non-stomatal control. Without this foundation, it is impossible to describe how *L*_p_ behaves across changing environments or to incorporate its effects into predictive frameworks.

In this study, we demonstrate that *L*_p_ is not a fixed property, but an active trait that responds predictably to changes in *ψ*_cy_. Through simultaneous measurements of gas exchange, *ψ*_cy_, and substomatal humidity, we show that the deployment of non-stomatal control of transpiration tracks closely with the decline in *ψ*_cy_ during transpiration. This mechanistic link positions *ψ*_cy_ as the key physiological signal driving *L*_p_, enabling internal regulation of water loss independent of stomatal aperture. Building on this relationship, we introduce a predictive model (Eq. (1)) that describes the shape and sensitivity of *L*_p_ engagement as a function of *ψ*_cy_. The model follows a logistic growth-type sigmoidal form, alike those commonly used in leaf hydraulic conductivity and vulnerability analyses^20,21^, providing a physiologically interpretable framework to quantify non-stomatal control of transpiration under varying internal water status (full derivation SI1).

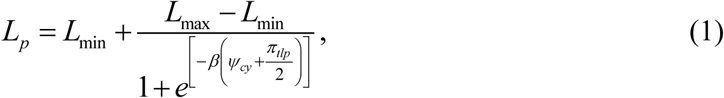

where *L*_min_ represents the minimum conductance (approximated to 0.33 mmol m^-2^ s^-1^ MPa^-1^ from our data), *L*_max_ is the maximum expected *L*_p_ (fitted to 7.69 mmol m^-2^ s^-1^ MPa^-1^ from our data), *π*_tlp_ is the osmotic pressure at the turgor loss point, and *β* is a positive parameter describing how responsive *L*_p_ is to changes in *ψ*_cy_ (*β* = *a*/*π*_tlp_, where *a* is a fitted dimensionless parameter).

Predicting *L*_p_ based on *ψ*_cy_ enables us to infer additional physiological variables that are otherwise inaccessible. By rearranging the water flow equation for the plasma membrane^7^, we estimate the water potential at the evaporative surface of the cell wall (*ψ*_wall_) from the difference between *ψ*_cy_ and the stomatal transpiration rate (*E*_s_) across *L*_p_ (Eq. (2)).

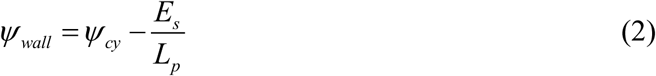

Joining this estimate with Kelvin’s equation in combination with Jurin’s Law, we derive the expected water vapour content in the substomatal cavity (*w*_i_) in equilibrium with *ψ*_wall_, which in turn determines the degree of substomatal unsaturation (Eq. (3)).

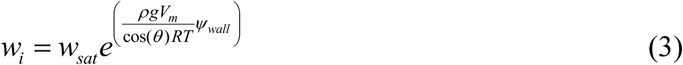

where *θ* is the contact angle of the water on wall fibrils (about 50° when it is wet; Kohonen ^22^), *R* the gas constant (8.314463 kg m^2^ s^-2^ °K^-1^ mol^-1^), *T* is the temperature (°K), *ρ* is the water mass density (1000 kg m^-3^), *g* is the gravitational acceleration (9.81 m s^-2^), and *V*_m_ the molar volume of the water (0.000018 m^3^ mol^-1^).

Together, equations (1) to (3) link measurable physiological variables to internal hydraulic conditions, allowing a continuous, model-based estimation of *w*_i_, and thus linking leaf hydraulics with gas exchange.

## 3 Results

The evolution of substomatal cavity unsaturation, transpiration rate, and stomatal conductance with increasing VPD for each evaluated leaf is shown in Figure 1. While each leaf exhibits a clear trend of decreasing relative humidity in the substomatal cavity as VPD increases (Figure 1a), the specific trajectory of unsaturation varies between leaves, showing no consistent pattern. A similar lack of consistency is observed for transpiration rate and stomatal conductance. Interestingly, *w*_i_/*w*_sat_ and *g*_sw_ often appear to follow antagonistic patterns—when one responds to increasing VPD, the other tends to remain unresponsive or changes with a much lower magnitude. Notably, if VPD is kept constant for long periods (40 to 60 minutes), stomatal conductance and non-stomatal control of transpiration adjust over time, usually towards a lower transpiration rate by either reducing stomatal conductance or *w*_i_. This observation aligns with the notion that these are independent mechanisms limiting water loss. However, the dynamics of this interaction are not easily deduced from simple comparisons of *w*_i_/*w*_sat_ and *g*_sw_, as their responses to VPD and *E* vary in direction and magnitude across leaves.

**Figure 1.**
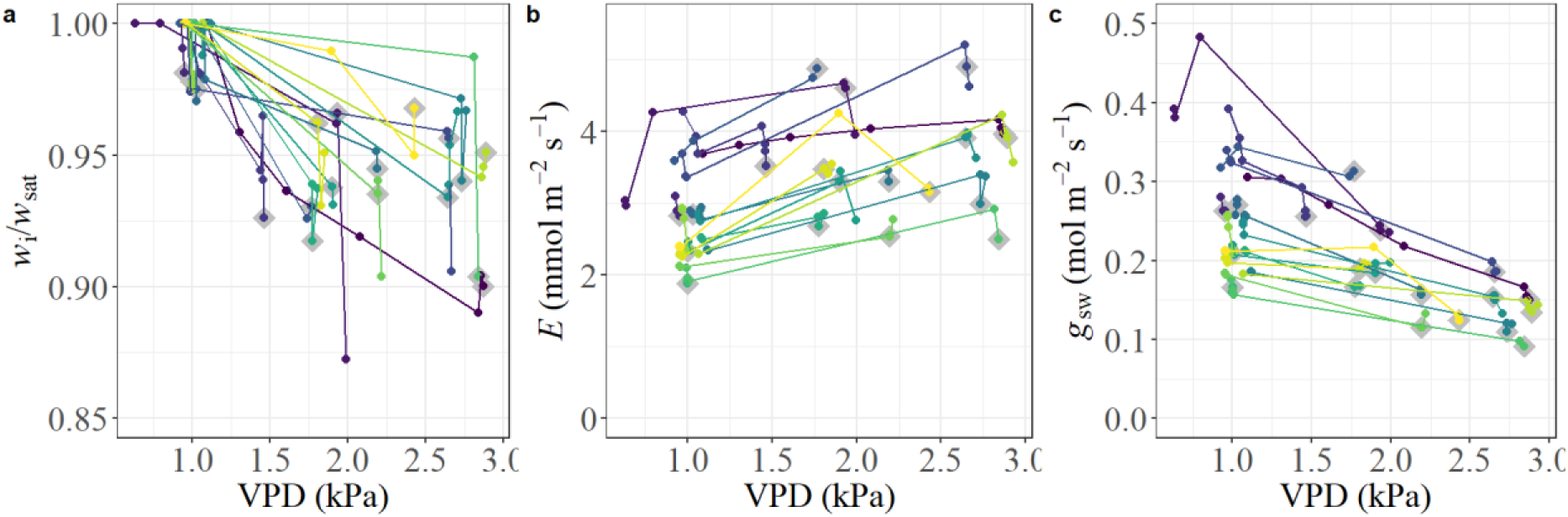
Leaf gas exchange responses to changes in vapour pressure deficit (VPD) in individual leaves. Each line represents a single leaf (19 colour-coded), with measurements taken after gas exchange stabilised. When VPD was changed, stability was reached after 15 to 50 minutes (variable across cases); when VPD was held constant and multiple measurements were taken, points were recorded at intervals of 20 to 30 minutes. Grey diamonds indicate the final time point on each leaf, when destructive sampling was conducted for cytosolic water potential (*ψ*cy) measurements. (a) Relative humidity in the substomatal cavity (*w*i/*w*sat) as a function of VPD. (b) Transpiration rate (*E*) as a function of VPD. (c) Stomatal conductance to water vapour (*g*sw) as a function of VPD.

The final measurement of the unsaturation versus VPD curve (Grey diamonds in Figure 1) represents the moment when samples were collected for calculating non-stomatal control of transpiration (*L*_p_) through destructive sampling. This set of samples provided a range of *w*_i_/*w*_sat_, *E*, and *r*_p_ values, allowing us to evaluate their correspondence with one another.

In summary of the findings presented below, we observed a clear correlation between the deployment of non-stomatal control of transpiration (*L*_p_) and the decline in cytosolic water potential (*ψ*_cy_), as well as with the progression of internal unsaturation. By contrast, we found no consistent relationship between *L*_p_ changes and either transpiration rate (*E*) or vapour pressure deficit (VPD). These findings are explored in detail in the following sections.

### 3.1 Non-stomatal control of transpiration and cytosolic water potential

Our results show that the non-stomatal control of transpiration, *L*_p_, followed a clear and consistent pattern of deployment as *ψ*_cy_ declined (^F^igure ^2^a). The decline in *ψ*_cy_ also aligned closely with the progression of unsaturation in the substomatal cavity (Figure 2b). Notably, the response of *L*_p_ to *ψ*_cy_ was not uniform: in the early stages of *ψ*_cy_ decline, *L*_p_ drops at a relatively stable pace. However, once a threshold in *ψ*_cy_ was crossed (about -1.2 MPa in our data), *L*_p_ decreases sharply and rapidly plateaus towards its minimum, suggesting a non-linear or sigmoidal relationship. This behaviour was mirrored by the evolution of unsaturation, where *w*_i_/*w*_sat_ gradually decreased in parallel with the drop in *L*_p_.

**Figure 2.**
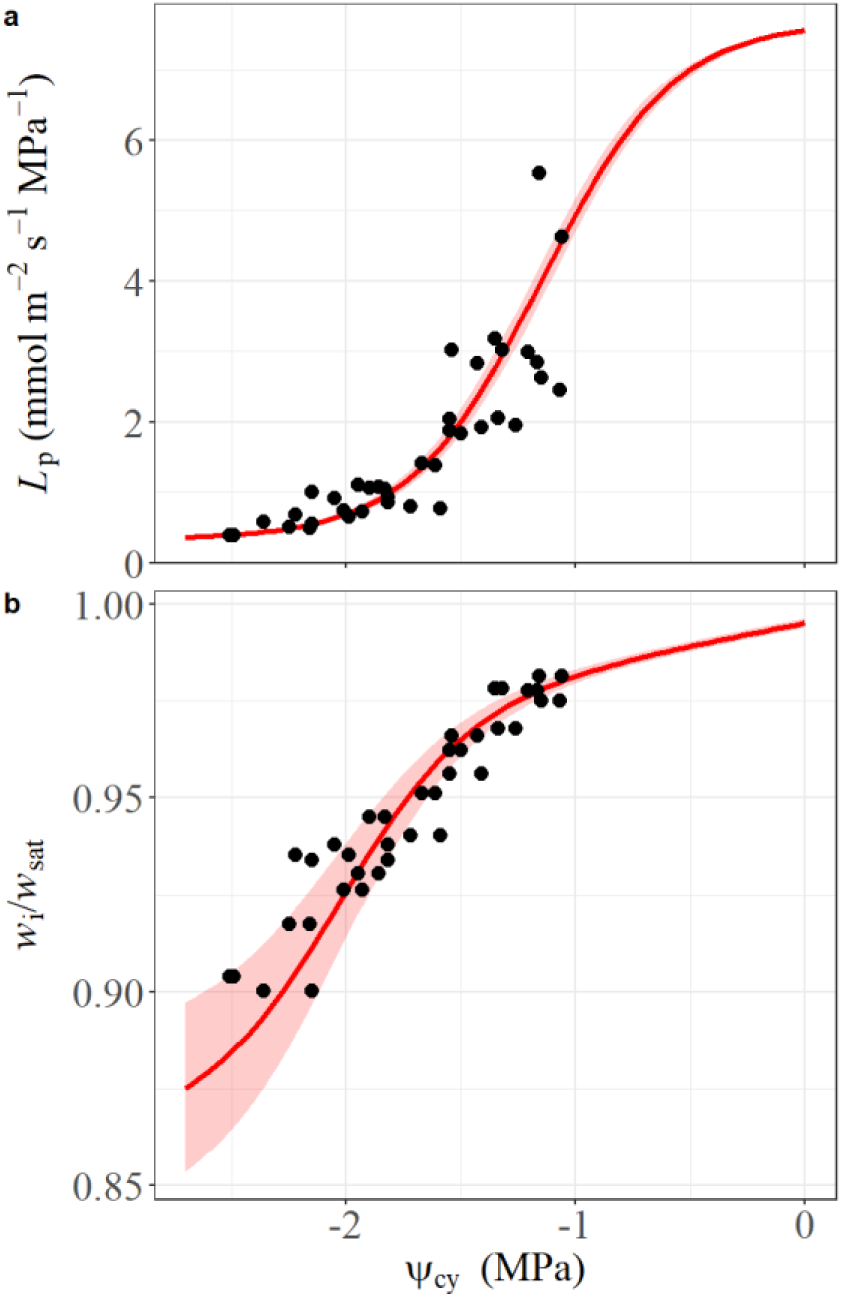
Correlation between cytosolic water potential, substomatal cavity unsaturation, and the non-stomatal control of transpiration (*L*p). (a) Modelled and observed *L*p fitted using Equation 1. Shaded area represents the effect of variation in the turgor loss point (-2.3 ±0.08 MPa; SI2). Fitted parameters: *L*min = 0.33, *L*max = 7.69, *ψ*tlp = -2.3, *β* = 3.5. (b) Modelled and observed *w*i/*w*sat, estimated from Equations 2 and 3, using modelled *L*p and an average transpiration rate of 3.3 mmol m^-2^ s^-1^. The shaded area represents the effect of variation in transpiration rate (3.3 ±0.8 mmol m^-2^ s^-1^).

As *L*_p_ decreased, *w*_i_/*w*_sat_ declined accordingly, consistent with restriction to water release to the cell wall and reduced vapour concentration in the airspace. Although the drop in *w*_i_/*w*_sat_ was less abrupt than the decrease in *L*_p_, both trends followed the same overall progression. This pattern suggests that changes in *L*_p_—driven by declining *ψ*_cy_—are a primary cause of airspace unsaturation under transpiring conditions rather than a direct response to VPD itself.

In Figure 2a, the red line shows the modelled evolution of *L*_p_ as a function of *ψ*_cy_, based on Equation (1). This curve was generated using a measured value of *π*_tlp_ (from pressure–volume analysis; see SI2), with *L*_min_ estimated from observed conductance under high VPD conditions in our data, and *L*_max_ and *β* fitted to the full dataset. The shaded area represents a sensitivity analysis reflecting the effect of variation in *π*_tlp_ among our samples. In Figure 2b, the red line corresponds to the predicted *w*_i_/*w*_sat_ values obtained by applying Equations (2) and (3), using *L*_p_ values from Equation (1) and the average *E* observed across all measurements, which was 3.3 mmol m^-2^ s^-1^. The shaded area in this panel represents a sensitivity analysis showing how the observed variation in *E* (±0.8 mmol m^-2^ s^-1^) affects the predicted degree of substomatal cavity unsaturation.

As expected from our modelling (Equations (1) to (3)), the predicted relationship between *L*_p_ and *w*_i_/*w*_sat_ reflects an asymptotic trend (SI3 Fig. S2). Relative humidity in the substomatal cavity plateaus asymptotically towards saturation at high *L*_p_, but declines sharply with decreasing *L*_p_, consistent with physical constraints on water movement toward the cell wall.

### 3.2 Modelling physiological constraints on *L*_p_ behaviour

To describe the relationship between *ψ*_cy_ and the deployment of *L*_p_, we used a logistic growth-type function (Eq. (1); Figure 2a). This choice was guided by the observed pattern in our data: *L*_p_ dropped steadily at high *ψ*_cy_, but decreased rapidly as *ψ*_cy_ declined and approached the osmotic pressure at turgor loss point (*π*_tlp_). This sigmoidal response is well captured by the flexibility of the logistic function, which allows for a sharp increase in resistance around a physiologically meaningful threshold (See derivation in SI1).

The function includes two constrained parameters that are also physiologically interpretable, *L*_min_ and *L*_max_. The minimum conductance value, *L*_min_, was derived from our data, based on the flat region in the curve at *ψ*_cy_ lower than -1.5 MPa. This value represents the baseline membrane conductance when undergoing dehydration. *L*_min_, defines the asymptotic lower bound of *L*_p_ as water stress increases. We set this parameter to be 0.33 mmol m^-2^ s^-1^ MPa^-1^. This value was directly observed and projected based on the trajectory of *L*_p_ under declining *ψ*_cy_ and is consistent with the rapid decrease in conductance near *π*_tlp_ (Figure 2a).

*L*_max_ was fitted, but from a physiological perspective, the magnitude of *L*_max_ can be constrained by considering the typical *ψ*_cy_ in well-hydrated leaves. Experimental measurements place *ψ*_cy_ near -0.3 MPa under full turgor conditions^23^, which implies that, to maintain high relative humidity in the substomatal cavity even under mild transpiration, membrane conductance must be sufficiently high. This sets a realistic lower bound on *L*_max_ in the modelling framework.

In standard pressure-volume curve analyses, full turgor is often assigned a water potential of 0 MPa for simplicity^24^, but this assumption would imply that either *L*_max_ is infinite or, worse, that negative conductance would be required to maintain *ψ*_cy_ below 0. Our estimated *L*_max_ of approximately 7.69 mmol m^-2^ s^-1^ MPa^-1^ at low *E* corresponds to a *ψ*_cy_ of -0.3 MPa and results in a substomatal relative humidity near 99%, consistent with the absence of unsaturation in fully hydrated-unstressed leaves. Note that significantly lower values of *L*_max_ would lead to noticeably lower *w*_i_/*w*_sat_ even at low transpiration rates, which would be easily detectable in standard gas exchange measurements but are not observed. Thus, while *L*_max_ was fitted from data, its value is supported both by physical reasoning and by observed physiological baselines.

The function is anchored on the difference *ψ*_cy_ - *π*_tlp_/2, positioning *π*_tlp_/2 at the centre of the logistic transition. As *ψ*_cy_ approaches -*π*_tlp_, *L*_p_ decreases steeply until it finds a plateau. At high *ψ*_cy_, when cells are fully turgid, *L*_p_ is maximal; as water potential declines, resistance rises sharply, contributing to reduced water efflux and stabilisation of internal water status. The parameter *β* describes the *L*_p_ engagement sensitivity, a trait reflecting how strongly a given species or condition regulates *L*_p_ in response to internal water status.

This pattern is related to traditional vulnerability curves used in leaf hydraulics, where hydraulic conductivity declines with decreasing water potential^25^. However, those curves usually describe the composite conductivity of the whole leaf, including the xylem^26^, while we focus specifically on the mesophyll membrane. In the range between full turgor and *π*_tlp_, our data suggest that *L*_p_ is the dominant component of total resistance. This must be the case, as turgor is maintained, and any greater downstream resistance would have resulted in further cytosolic dehydration. Beyond *π*_tlp_, other resistances—such as those in the xylem—may become more limiting, but these fall outside the range captured in our measurements (Figure 2a).

The logistic function provides a physiologically grounded and empirically supported framework to model changes in *L*_p_. It captures the observed non-linear decrease in membrane conductance as a function of cytosolic water potential and allows for the integration of non-stomatal regulation into leaf hydraulic and gas exchange models.

### 3.3 Non-stomatal control deployment does not follow transpiration demand evolution

The deployment of non-stomatal control, reflected in changes in *L*_p_, did not follow any consistent pattern with either transpiration rate or VPD (Figure 3a and b). Similarly, the degree of substomatal unsaturation (*w*_i_/*w*_sat_) also showed no clear dependence on these drivers alone (Figure 3c and d). This reinforces the interpretation that neither VPD nor transpiration rate is, by itself, the physiological trigger for a decrease in *L*_p_. Rather, transpiration appears to respond to the decrease in *L*_p_, not the other way around. This further supports the role of *L*_p_ as a true non-stomatal control of transpiration, governed by internal water status rather than ambient atmospheric conditions.

**Figure 3.**
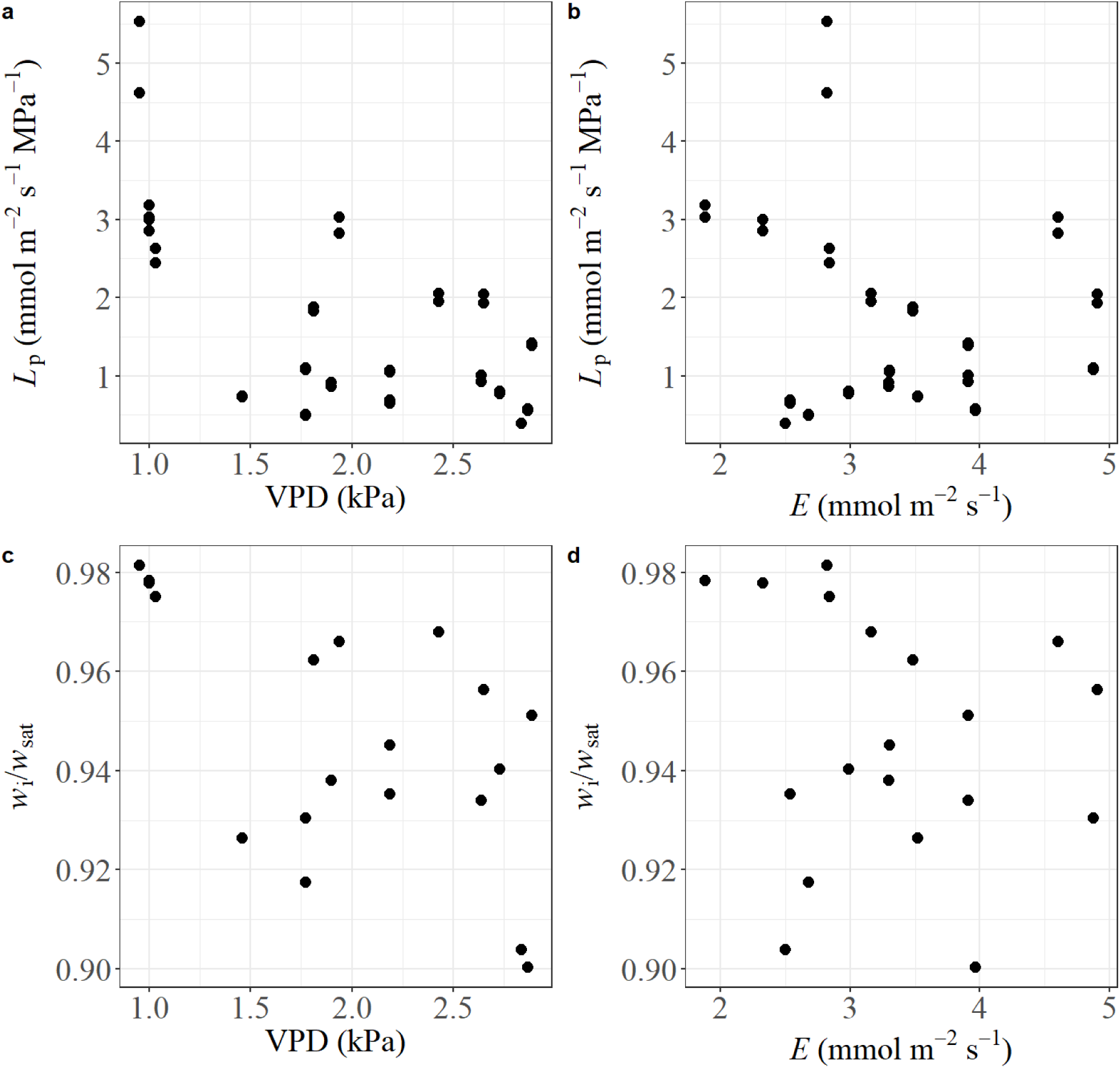
Relationships between the non-stomatal control of transpiration (*L*p), substomatal cavity unsaturation (*w*i/*w*sat), and key gas exchange drivers. (a, b) *L*p as a function of vapour pressure deficit (VPD) and transpiration rate (*E*), respectively. (c, d) *w*i/*w*sat plotted against VPD and *E*, respectively. All values were derived from combined gas exchange and water potential measurements obtained at the time of the destructive sampling indicated by grey diamonds in Figure 1.

While our data in Figure 3 were obtained from different leaves, and individual physiological variability may obscure subtle trends, the observed patterns support the view that non-stomatal regulation is not mediated by transpiration or VPD. This stands in contrast with the behaviour observed for stomatal control, where stomata have been shown to respond to transpiration rate^27^, implying a feedback mechanism based on water loss. In our case, however, the non-stomatal control of transpiration shows no such relationship. Instead, *L*_p_ responds to internal water status, independent of VPD and transpiration rate. This distinction reinforces the notion that non-stomatal regulation may operate via a fundamentally different physiological trigger than stomatal control, not as a response to flux but as a driver of it.

Moreover, when comparing *L*_p_ and stomatal conductance across leaves, no consistent coordination emerges (Figure 4). The absence of a clear relationship between *L*_p_ and *g*_sw_ lends further support for the interpretation that *L*_p_ deployment does not simply scale with stomatal conductance. Instead, it suggests that the two operate through, at least partially, distinct regulatory pathways, serving complementary roles under transpiring conditions.

**Figure 4.**
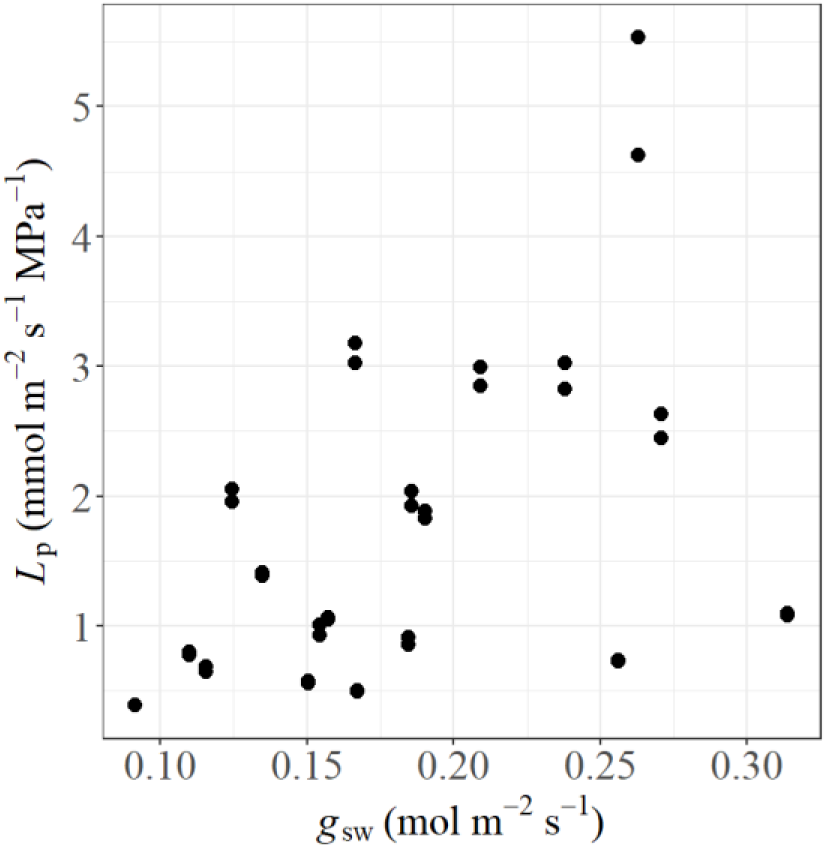
Relationship between non-stomatal control of transpiration (*L*p) and stomatal conductance (*g*sw) across sampled leaves. Each point represents a single leaf measurement taken at the time of cytosolic water potential sampling.

## 4 Discussion

Other studies have proposed that non-stomatal control of transpiration must involve an internal conductance that changes dynamically, located at the level of the mesophyll cell membranes^5^,^7,9,14,15^. Notably, Wong, et al. ^7^ observed a potential decline in *ψ*_cy_ during measurements of non-stomatal control of transpiration engagement. However, this variability was not directly and consistently quantified. Our results provide the first direct, point-by-point demonstration of systematic variation of *L*_p_, tightly aligned with the evolution of substomatal cavity unsaturation across multiple leaves (Figure 2). The consistency of this behaviour within and across individuals provides compelling evidence for a responsive, physiologically meaningful regulation of *L*_p_ in response to internal water status.

The strong and consistent relationship between *ψ*_cy_ and *L*_p_ reinforces *ψ*_cy_ as the likely mechanistic signal governing the non-stomatal control (Figure 2a). From a physiological perspective, this makes sense: a cell-based signal offers finer-scale responsiveness and some independence from stomatal cues. The capacity to modulate water loss based on internal water status, rather than relying on stomatal feedback, aligns with recent observations showing improved carbon gain under conditions where non-stomatal control is active^7,9^. This suggests that *L*_p_ adjustments contribute to maintaining water-use efficiency under stress without duplicating stomatal responses.

Previous studies speculated about possible triggers for non-stomatal control, including transpiration rate, leaf turgor status, and vapour pressure deficit (VPD)^7,9,14,17^. Turgor and transpiration have been the leading candidates, while VPD has occasionally been proposed due to its role in driving evaporative demand. In many experimental setups, VPD has been used to manipulate leaf water loss and provoke unsaturation^14^, which may have contributed to the perception of a direct link. However, our data show no consistent correlation between VPD or transpiration rate and the deployment of non-stomatal control (Figure 1), either in terms of *L*_p_ or substomatal unsaturation (Figure 3). Instead, these variables appear to act indirectly by triggering internal dehydration. By contrast, *ψ*_cy_ consistently explained the timing and degree of *L*_p_ decrease, clearly positioning it as the key physiological signal driving the response. It is tempting to speculate that the decrease in *L*_p_ is related to abscisic acid (ABA) production by mesophyll cells losing turgor, as ABA production has been shown to increase as leaf water potential declines^6,28^. However, this would require a physiological explanation for why ABA does not appear to affect stomata in an equivalent manner, given that no correlation was found between *L*_p_ and *g*_sw_ (Figure 4).

There has been a growing call in the literature for a mechanistic model that can describe how non-stomatal control of transpiration operates dynamically during transpiration^14,18,19^. Our approach addresses this gap by introducing a physiologically grounded function that links *L*_p_ to *ψ*_cy_ (Eq. (1)). The shape of the response follows a sigmoidal pattern well known in plant hydraulics, particularly from vulnerability curves describing xylem or leaf hydraulic conductance^26,29^. This correspondence was not imposed arbitrarily—it emerged from the data and lends physiological meaning to the model (See SI1). Our formulation provides a testable, quantitative framework that captures the non-linear nature of *L*_p_ modulation in relation to declining internal water status.

Because *ψ*_cy_ and xylem water potential are expected to remain close under most conditions— due to the lack of substantial resistance between the two compartments^30^—it is reasonable to speculate that leaf water potential measurements could be used as a proxy for our *L*_p_ model instead of *ψ*_cy_. This would allow the implementation of our approach in broader physiological models without requiring direct *ψ*_cy_ measurements. The resemblance between the *L*_p_–*ψ*_cy_ relationship and conventional *K*_leaf_–*ψ* relationships further supports the potential for generalisation and integration into existing hydraulic frameworks, though this will require further testing.

Finally, the introduction of the sensitivity parameter *β* opens the door to new questions about interspecific and intraspecific variation in non-stomatal control. While *π*_tlp_ sets the threshold, *β* defines how steeply resistance increases once that threshold is approached. It remains unclear whether *β* is relatively conserved across species or varies in response to ecological strategy, growth conditions, or evolutionary adaptation. Testing this across a broader range of species and environments will be an important step toward understanding the ecological relevance and plasticity of this control mechanism.

Several hypotheses could be proposed regarding the factors influencing variation in *β*. For example, species adapted to arid or high-VPD environments might evolve steeper (larger) *β* values, enabling a sharper and more decisive reduction in *L*_p_ once dehydration begins. Alternatively, *β* might reflect structural properties such as membrane composition, aquaporin density and gating behaviour, or mesophyll cell wall architecture. It is also possible that *β* is a plastic trait, modifiable in response to repeated stress exposure or developmental stage, potentially through acclimation or epigenetic regulation. Differences between photosynthetic lineages (e.g., C_3_ vs C_4_) may also play a role: previous work has shown that C_4_ species engage in non-stomatal control strongly under drying conditions^9^. It is plausible that they deploy this mechanism earlier in the dehydration process, potentially reflected in higher *ψ*_cy_ thresholds and distinct *β* values. As such, *β* may offer a powerful comparative tool to investigate the diversity, plasticity, and evolutionary significance of non-stomatal control across species and environments.

Non-stomatal control of transpiration has been observed in vascular plants^5–7,9^ and, more recently, an analogous—potentially ancestral—mechanism has been reported in bryophytes^31^, exhibiting similar functionality through conductance at the plasma membrane (i.e., *L*_p_). Non-stomatal control has been shown to limit transpiration without compromising carbon assimilation^9^, challenging long-standing assumptions about the trade-off between water conservation and photosynthetic gain. In this study, we provide the first quantitative evidence that the deployment of non-stomatal control is linked to internal water status and operates independently of stomatal behaviour. This mechanistic insight offers a new framework for understanding how plants regulate water loss at the cellular level.

Important questions remain. If non-stomatal control can reduce water loss without limiting CO_2_ uptake, why is it not engaged continuously? What physiological or ecological trade-offs constrain its deployment? How does it interact with other transpiration-dependent processes, such as thermal regulation or nutrient transport? While these questions remain, our findings lay a foundation for addressing them. By providing a predictive model of non-stomatal control dynamics and identifying cytosolic water potential as a key regulatory signal, we take a critical step toward integrating this mechanism into the broader framework of plant water relations. Future work exploring its variability, coordination with stomatal regulation, and implications across environments will be key to unlocking its full significance.

## 5 Conclusions

Our results demonstrate that mesophyll membrane conductance to water flow (*L*_p_) is not static, but dynamically adjusted during transpiration. This adjustment is closely and consistently linked to cytosolic water potential (*ψ*_cy_), identifying *ψ*_cy_ as the most likely mechanistic signal underlying the deployment of non-stomatal control of transpiration. We showed that neither transpiration rate nor vapour pressure deficit (VPD) explained the observed *L*_p_ changes, further reinforcing the central role of internal water status in regulating this process.

To formalise this relationship, we developed a mechanistic model in which *L*_p_ decreases nonlinearly as *ψ*_cy_ declines. The shape of this response resembles well-established patterns from leaf hydraulic vulnerability curves, offering a physiologically interpretable framework, linking hydraulics with gas exchange. Together, these findings advance our understanding of how plants regulate water loss beyond the stomatal pathway, highlighting a dynamic, internal layer of control that operates in parallel with external environmental cues.

## 6 Materials and methods

### 6.1 Plant material

We conducted experiments on nineteen leaves harvested from six cotton plants (*Gossypium hirsutum*), with three leaves sampled from each plant, except for one plant from which four were taken. These plants were grown from seeds in ten-litre pots filled with Martins Potting Mix (sourced from Martins Fertilizers, Yass, NSW, Australia). When planting, 5 grams of Osmocote Exact slow-release fertiliser (provided by Scotts Australia, Bella Vista, NSW, Australia) were incorporated into the soil. The plants were maintained in a glasshouse where natural sunlight was the primary light source. A controlled temperature regime of 28°C by day and 20°C by night was applied. Daily irrigation was carried out to meet the plants’ water requirements. Our investigation focused on leaves that had reached full maturity.

The total cuticular conductances to water vapour for the leaves were estimated through the Red-light technique, as introduced by Márquez, et al. ^32^, utilising an LI-6800 (LI-COR, Lincoln, NE, USA). The measurements yielded conductances within a 4.4±0.6 mmol m^-2^ s^-1^ range, denoted as an average with standard deviation. It was assumed that the cuticular conductance to CO_2_ is 5% of that to water vapour^33^.

### 6.2 Equipment

Experiments were conducted using two LI-6800 gas exchange analysis systems (LI-COR, Lincoln, NE, USA) connected, as shown in ^34^, to independently measure adaxial and abaxial gas exchange. Additional experiments were conducted using an LI-6800 with a 6 cm^2^ chamber (6800-01A). The gas exchange parameters were determined based on equations presented by Márquez, et al. ^33^.

Water potential measurements were performed using two leaf psychrometers PSY1 (ICT International, Australia).

### 6.3 Turgor loss point determination

Pressure-volume (PV) curves were generated to characterise the water relations of leaves from six individual plants (one leaf per plant). Measurements of leaf water potential (*ψ*_leaf_) and corresponding fresh mass were taken repeatedly as leaves dehydrated gradually.

To determine *ψ*_leaf_, a Scholander-type pressure chamber was employed. Leaves were harvested during the night. Prior to excision, petioles were immersed in water to prevent xylem embolism during detachment, a second cut under the water was performed about half a centimetre from the original cut. Immediately after collection, leaves were placed in a dark chamber at 5°C for 12 hours, with their petioles submerged in water to allow full rehydration. Following rehydration, leaves were transferred to ambient laboratory conditions (c. 23 °C) and acclimatised for 45 minutes in a sealed plastic bag to maintain high humidity and prevent water loss.

Fresh mass at full turgor was recorded for each leaf. Subsequently, leaves were allowed to dry naturally at room temperature. At regular intervals, leaves were removed and weighed, and their water potential measured using the pressure chamber. This process was repeated ten times per leaf, spanning a range from full turgor to approximately -3 MPa.

After the final water potential measurement, all leaves were oven-dried at 75°C to constant weight in order to determine dry mass. Gravimetric water content at each time point was then calculated as the difference between fresh mass and dry mass. Pressure-volume curves were constructed by plotting -1/*ψ*_leaf_ against relative water content. The turgor loss point (*ψ*_tlp_) was determined visually as the point of inflexion where the curve transitions from a linear to a non-linear phase, following standard procedures.

### 6.4 Unsaturation Measurements

The gas exchange measuring sequence started at low vapour pressure deficit (VPD), around 0.7 to 0.8 kPa, and *c*_a_ for both upper and lower leaf cuvettes at 400 µmol mol^-1^. Light intensity was set at 1000 µmol m^-2^ s^-1^. Once steady-state gas exchange rates were achieved, the lower cuvette’s CO_2_ concentration (*c*_a_) was decreased to achieve an assimilation rate on the lower leaf surface of 0 ±0.3 μmol m^-2^ s^-1^ (zero).

The method for determining the internal air space resistance to CO_2_ diffusion (*R*_ias_) was adapted from the procedure by Wong, et al. ^7^. This required sustaining an atmospheric CO_2_ concentration in the upper cuvette at 400 µmol mol^-1^, while the CO_2_ concentration in the lower cuvette is decreased until the assimilation rate of the lower surface is zero. Under these conditions, the CO_2_ concentration within the substomatal cavity (*c*_i_) on the abaxial surface equalled the leaf’s minimal CO_2_ concentration. Under this premise, Eq. (4) was utilised to calculate *R*_ias_, adopting a value of m = -0.31, as Márquez, et al. ^8^ proposed.

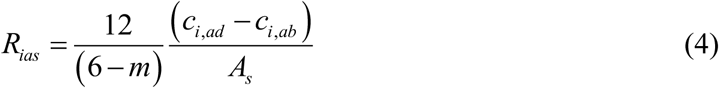

At the lower VPD, the substomatal vapour concentration (*w*_i_) was assumed to equal the saturated vapour concentration at leaf temperature (*w*_sat_). VPD increases were performed from ∼0.8 kPa to different values between 1 and 2.9 kPa, aiming to achieve different levels of unsaturation. The VPD increase was performed in a linear increment of 10 minutes duration. After achieving unsaturation and constant gas exchange parameters at the target VPD, usually taking 20 to 30 minutes, the leaf was removed from the chamber, and two leaf discs were taken with a 6 mm leaf punch and placed in the psychrometer chamber. Prominent veins were avoided when taking leaf punches.

To estimate unsaturation, Eq. (5) presented by Márquez, et al. ^9^ was used to compute the actual *w*_i_ assuming constant *R*_ias_; the positive root was taken.

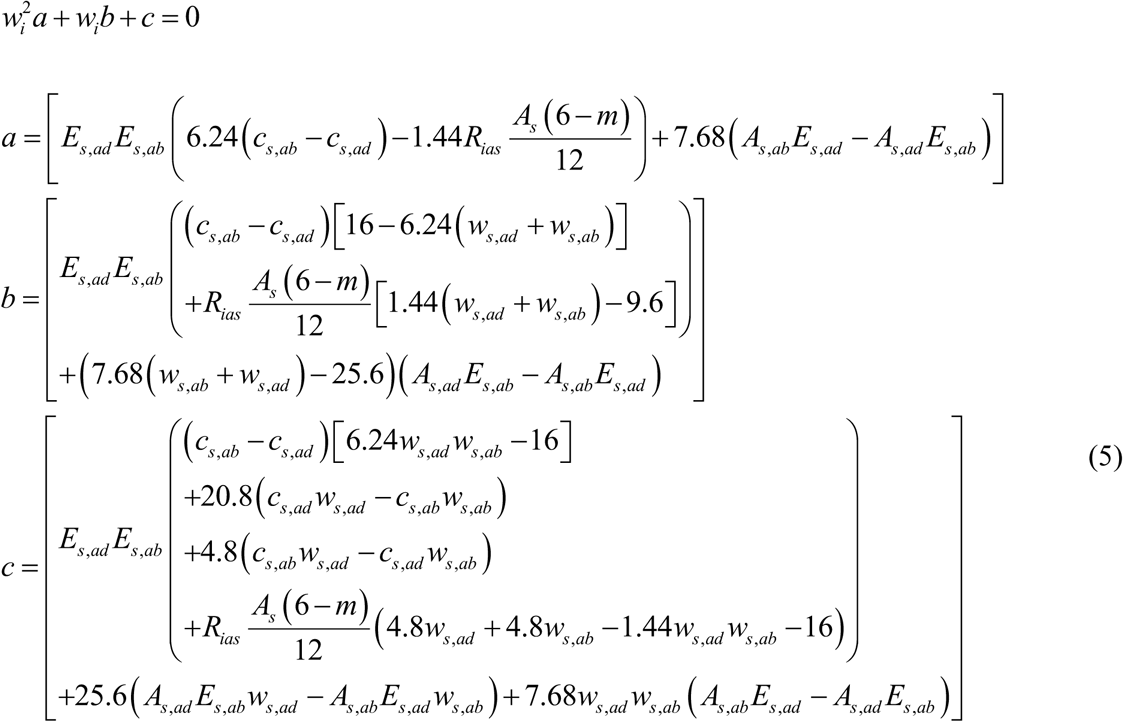

where *E*_s_ is the transpiration rate through the stomata, *c*_s_ is CO_2_ concentration at the leaf surface, *A*_s_ is the assimilation rate through the stomata, *w*_s_ is the vapour concentration at the leaf surface, and *ad* and *ab* subscripts refer to the adaxial and abaxial surface, respectively.

Then, relative humidity in the substomatal cavity was estimated as *w*_i_/*w*_sat_.

### 6.5 Cell membrane conductance (*L*_p_) measurements

We define *L*_p_ as the apparent resistance to water flow across mesophyll cell membranes, as inferred from gas exchange measurements in a projected leaf area. While *L*_p_ does not represent the permeability of individual membranes, it reflects a bulk property that emerges from the composite effect of many membranes under transpiring conditions. As shown by Wong, et al. ^7^, this resistance is physically located in the plasma membrane–cell wall interface, and their estimates fall within known physiological ranges. Thus, although *L*_p_ is observed at the tissue level, its origin can be confidently attributed to changes in plasma membrane conductivity, so that

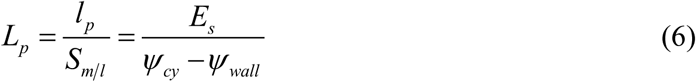

where *l*_p_ is the resistance to water in the plasma membrane, and *S*_m/l_ is the ratio of mesophyll surface over projected leaf surface area.

Psychrometer measurements were interpreted as cytosolic water potential (*ψ*_cy_), as the cytosol and vacuole together contain the vast majority of water in the leaf.

We can estimate the water potential in the liquid phase of the cell wall (*ψ*_wall_) from the vapour pressure equivalent in equilibrium in the substomatal cavity using the Kelvin equation in combination with Jurin’s law (Eq. (7)):

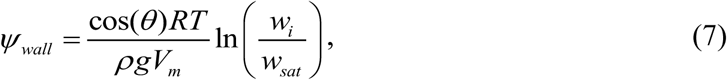

where *θ* is the contact angle of the water on wall fibrils (about 50° when is wet; Kohonen ^22^), *R* the gas constant (8.314463 kg m^2^ s^-2^ °K^-1^ mol^-1^), *T* is the temperature (°K), *ρ* is the water mass density (1000 kg m^-3^), *g* gravitational acceleration (9.81 m s^-2^), *V*_m_ the molar volume of the water (0.000018 m^3^ mol^-1^) and *w*_i_/*w*_sat_ is the relative humidity in equilibrium with the liquid phase.

## 7 Acknowledgements

DAM acknowledges the Pump Priming Award from the School of Biosciences, University of Birmingham, which supported equipment acquisition. This work was also supported by the Australian Research Council (Discovery Grant DP210103186, awarded to GDF and LAC) and by the Natural Environment Research Council (grant NE/W00674X/1, awarded to FAB).

## 8 Author contributions

DAM and GDF developed the initial concept and the original formulation of the theoretical framework. DAM conducted the gas exchange experiments and water potential measurements. LAC and FAB support refining and elaborating the theoretical framework and data interpretation. All authors contributed to the writing and revision of the final version.

## Supplementary Information

### SI1 *L*_p_ model derivation

We propose that variation in *L*_p_ occurs in response to changes in cytosolic water potential (*ψ*_cy_), and that this variation scales proportionally with the ratio *ψ*_cy_/*π*_tlp_. This ratio allows us to express the rate of change of *L*_p_ in the following form:

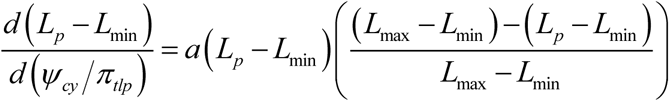

We expect *L*_p_ to be constrained between a maximum conductance (*L*_max_) and a minimum conductance (*L*_min_). These act as asymptotic limits: *L*_max_ when *ψ*_cy_ is close to full turgor (i.e., near 0 MPa), and *L*_min_ as *ψ*_cy_ becomes increasingly negative due to dehydration. The transition between these limits, and how abruptly it occurs across the physiological range, is controlled by a dimensionless steepness parameter *a*.

Note that we have preserved the structure of a logistic form, which makes it clear that this expression describes a standard logistic differential equation. For simplicity, we now define: *s*=*L*_p_-*L*_min_; *x*=*ψ*_cy_/*π*_tlp_; Δ*L*=*L*_max_-*L*_min_. This allows us to rewrite the equation as:

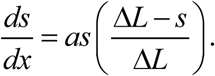

Solving this differential equation:

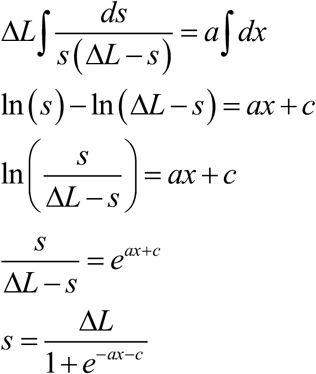

Returning to our original parameters, we obtain:

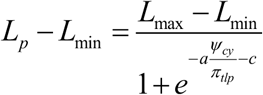

From our theoretical framework, we hypothesise that *L*_p_ operates functionally between full turgor (*ψ*_cy_ ≈ 0) and the turgor loss point (*π*_tlp_), and that the ratio *ψ*_cy_/*π*_tlp_ captures this physiological range in a normalised form. Within this interval, the logistic transition is expected to occur most rapidly at the midpoint of the range, that is, when *ψ*_cy_ ≈ *π*_tlp_/2. This suggests that the inflexion point of the sigmoidal response lies at *x* = 1/2, and therefore arises directly from our hypothesis, and is supported by our data. Incorporating this into our model corresponds to setting *c* = 1/2, which yields:

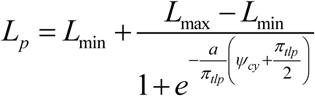

### SI2 Pressure-volume curves

Pressure-volume curves were obtained from leaves of six individual plants. The estimated turgor loss point (*ψ*_tlp_) ranged from -2.45 to -2.22 MPa, with a mean of -2.30 ± 0.08 MPa (Fig. S1). These values were consistent among replicates and fall within the upper range reported for the species. They were used to parameterise subsequent modelling of *r*_p_ and substomatal humidity.

**Fig. S1.**
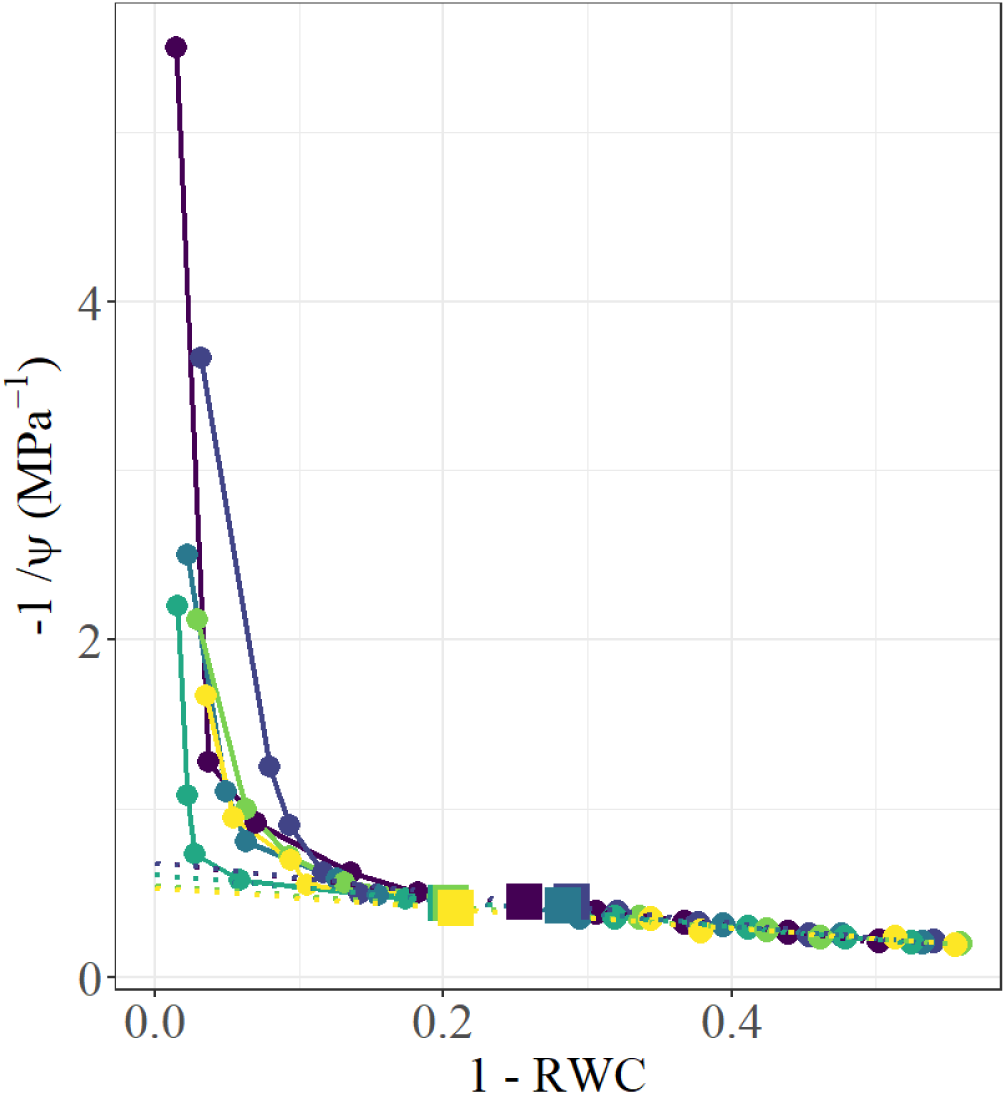
Water potential response to relative water content in leaves from six individual plants. Each colour represents a single leaf from a different plant, showing the inverse water potential (-1/ψ) as a function of relative water loss (1 – RWC). Solid lines represent measurements within the turgor range, while dotted lines show linear fits to the post-turgor phase. Square symbols indicate the turgor loss point (*ψ*tlp) for each leaf, plotted at the corresponding (1 – RWC, -1/*ψ*tlp) coordinate.

### SI3 relationship between *L*_p_ and *w*_i_/*w*_sat_

To illustrate how the relationship between mesophyll membrane conductance (*L*_p_) and substomatal humidity arises from our mechanistic framework, we combined Equations (1) to (3) from the main text. Substituting Equation (2) into Equation (3) yields an explicit expression for *w*_i_/*w*_sat_ as a function of *L*_p_:

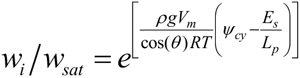

Neither *ψ*_cy_ nor *E*_s_ are constant in a physiological sense; both vary under transpiring conditions. However, in this formulation, the term *E*_s_/*L*_p_ dominates the shape of the response because it is expected to vary more strongly in magnitude than *ψ*_cy_ over the range of interest. The resulting relationship is asymptotic: at low *L*_p_, small increases in conductance cause large increases in *w*_i_/*w*_sat_, while at high *L*_p_, the response plateaus as saturation is approached. This behaviour is visualised in Fig. S2, where *w*_i_/*w*_sat_ is plotted as a function of *L*_p_, confirming the trend predicted by our model.

**Fig. S2.**
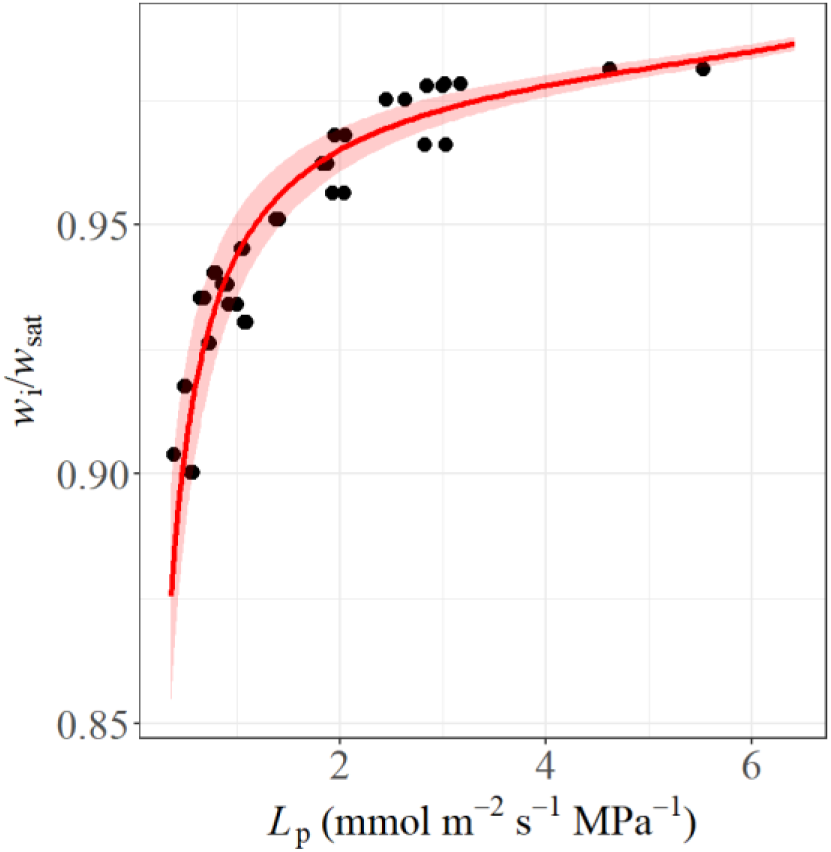
Modelled relationship between non-stomatal control of transpiration (*L*p) and relative humidity in the substomatal cavity (*w*i/*w*sat), based on Equations (1) to (3) from the main text. The red line represents the predicted mean *w*i/*w*sat as a function of *L*p, while the shaded area shows the variation arising from the observed range in transpiration rate (*E*s = 3.3 ± 0.8 mmol m^-2^ s^-1^). Black points represent observed values.

## References

1 Gaastra, P. *Photosynthesis of crop plants as influenced by light, carbon dioxide, temperature, and stomatal diffusion resistance*. (Meded. Landbouwhogeschool, Wageningen, 1959).

2 Farquhar, G. D. & Wong, S. C. An Empirical Model of Stomatal Conductance. Functional Plant Biology 11, 191–210 (1984). 10.1071/PP9840191

3 Buckley, T. N., Mott, K. A. & Farquhar, G. D. A hydromechanical and biochemical model of stomatal conductance. PC&E 26, 1767–1785 (2003). 10.1046/j.1365-3040.2003.01094.x

4 Lawson, T. & Matthews, J. Guard Cell Metabolism and Stomatal Function. Annual Review of Plant Biology 71, 273–302 (2020). 10.1146/annurev-arplant-050718-100251

5 Cernusak, L. A. et al. Unsaturation of vapour pressure inside leaves of two conifer species. Scientific Reports 8, 1–7 (2018). 10.1038/s41598-018-25838-2

6 Cernusak, L. A., Goldsmith, G. R., Arend, M. & Siegwolf, R. T. W. Effect of vapor pressure deficit on gas exchange in wild-type and abscisic acid–insensitive plants. Plant Physiology 181, 1573–1586 (2019). 10.1104/pp.19.00436

7 Wong, S. C. et al. Humidity gradients in the air spaces of leaves. Nat. Plants 8, 971–978 (2022). 10.1038/s41477-022-01202-1

8 Márquez, D. A., Stuart-Williams, H., Cernusak, L. A. & Farquhar, G. D. Assessing the CO_2_ concentration at the surface of photosynthetic mesophyll cells. New Phytol. 238, 1446–1460 (2023). 10.1111/nph.18784

9 Márquez, D. A., Wong, S. C., Stuart-Williams, H., Cernusak, L. A. & Farquhar, G. D. Mesophyll airspace unsaturation drives C_4_ plant success under vapour pressure deficit stress. PNAS 121 (2024). 10.1073/pnas.240223312

10 Jarvis, P. G. & Slatyer, R. O. The role of the mesophyll cell wall in leaf transpiration. Planta 90, 303–322 (1970). 10.1007/BF00386383

11 Farquhar, G. D. & Raschke, K. On the resistance to transpiration of the sites of evaporation within the leaf. Plant Physiology 61, 1000–1005 (1978). 10.1104/pp.61.6.1000

12 Canny, M. J. & Huang, C. X. Leaf water content and palisade cell size. New Phytol. 170, 75–85 (2006). 10.1111/j.1469-8137.2005.01633.x

13 Canny, M. Water loss from leaf mesophyll stripped of the epidermis. Functional Plant Biology 39, 421–434 (2012). 10.1071/FP11265

14. 14 Cernusak, L. A., et al. Unsaturation in the air spaces of leaves and its implications. PC&E (2024). 10.1111/pce.15001

15 Jain, P. et al. Localized measurements of water potential reveal large loss of conductance in living tissues of maize leaves. Plant Physiology (2023). 10.1093/plphys/kiad679

16 Jain, P. et al. New approaches to dissect leaf hydraulics reveal large gradients in living tissues of tomato leaves. New Phytol. 242, 453–465 (2024). 10.1111/nph.19585

17 Buckley, T. N. & Sack, L. The humidity inside leaves and why you should care: implications of unsaturation of leaf intercellular airspaces. American Journal of Botany 106, 618–621 (2019). 10.1002/ajb2.1282

18 Rockwell, F. E. et al. Extreme undersaturation in the intercellular airspace of leaves: a failure of Gaastra or Ohm? Annals of Botany 130, 301–316 (2022). 10.1093/aob/mcac094

19 Márquez, D. A., Gardner, A. & Busch, F. A. Navigating challenges in interpreting plant physiology responses through gas exchange results in stressed plants. Plant Ecophysiology (2025).

20 Cochard, H. Vulnerability of several conifers to air embolism. Tree Physiology 11, 73–83 (1992).

21 Cochard, H., Cruiziat, P. & Tyree, M. T. Use of Positive Pressures to Establish Vulnerability Curves 1: Further Support for the Air-Seeding Hypothesis and Implications for Pressure-Volume Analysis. Plant Physiology 100, 205–209 (1992). 10.1104/pp.100.1.205

22. 22 Kohonen, M. M. in Contact Angle, Wettability and Adhesion Vol. 5 47–57 (CRC Press, 2008).

23 Tyree, M. T. & Hammel, H. T. The Measurement of the Turgor Pressure and the Water Relations of Plants by the Pressure-bomb Technique. Journal of Experimental Botany 23, 267–282 (1972).

24 Bartlett, M. K., Scoffoni, C. & Sack, L. The determinants of leaf turgor loss point and prediction of drought tolerance of species and biomes: a global meta-analysis. Ecol. Lett. 15, 393–405 (2012). 10.1111/j.1461-0248.2012.01751.x

25 Scoffoni, C., McKnown, A. D., Rawls, M. & Sack, L. Dynamics of leaf hydraulic conductance with water status: quantification and analysis of species differences undes steady state. Journal of Experimental Botany 63, 643–658 (2012).

26 Scoffoni, C. et al. Leaf vein xylem conduit diameter influences susceptibility to embolism and hydraulic decline. New Phytol. 213, 1076–1092 (2017).

27 Mott, K. A. & Parkhurst, D. F. Stomatal responses to humidity in air and helox. PC&E 14, 509–515 (1991). 10.1111/j.1365-3040.1991.tb01521.x

28 Pierce, M. & Raschke, K. Correlation between loss of turgor and accumulation of abscisic acid in detached leaves. Planta 148, 174–182 (1980). 10.1007/BF00386419

29 Sack, L., Buckley, T. N. & Scoffoni, C. Why are leaves hydraulically vulnerable? Experimental botany 67, 4917–4919 (2016).

30 Scoffoni, C. et al. Outside-Xylem Vulnerability, Not Xylem Embolism, Controls Leaf Hydraulic Decline during Dehydration. Plant Physiology 173, 1197–1210 (2017). 10.1104/pp.16.01643

31 Perera-Castro, A. V., Márquez, D. A., Busch, F. A. & Hanson, D. Evidence for active regulation of transpiration in non-stomatal plants. bioRxiv, 2025.2004.2001.646543 (2025). 10.1101/2025.04.01.646543

32 Márquez, D. A., Stuart-Williams, H., Farquhar, G. D. & Busch, F. A. Cuticular conductance of adaxial and abaxial leaf surfaces and its relation to minimum leaf surface conductance. New Phytol. 233 (2022). 10.1111/nph.17588

33 Márquez, D. A., Stuart-Williams, H. & Farquhar, G. D. An improved theory for calculating leaf gas exchange more precisely accounting for small fluxes. Nat. Plants 7, 317–326 (2021). 10.1038/s41477-021-00861-w

34 Márquez, D. A., Stuart-Williams, H., Wong, S. C. & Farquhar, G. D. An improved system to measure leaf gas exchange on adaxial and abaxial surfaces. Bio-protocol 13, e4687 (2023). 10.21769/BioProtoc.4687

